# THC Reverses SIV-Induced Senescence in Astrocytes: Possible Compensatory Mechanism Against HIV Associated Brain Injury?

**DOI:** 10.1101/2025.05.16.654476

**Authors:** Alison R. Van Zandt, Miranda D. Horn, Tiffany A. Peterson, Sarah Y. Dickinson, Elise M. Frost, Andrew G. MacLean

## Abstract

Despite effective combination antiretroviral therapy (cART), chronic neuroinflammation and glial dysfunction continues to be an important yet understudied issue with people living with HIV (PLWH). The endocannabinoid system is increasingly recognized as a potential therapeutic target for modulating neuroimmune environments, given its role in regulating synaptic plasticity, immune responses, and neuroinflammatory cascades. However, the extent to which cannabinoids influence HIV-associated neuroinflammation remains unclear. This study investigates the impact of Δ9-tetrahydrocannabinol (THC) on astrocyte growth characteristics, viability, and senescence-associated cytokine release following exposure to HIV Tat protein using primary mixed glial cultures derived from rhesus macaques. Real-time impedance-based cellular integrity assessments were conducted using the xCELLigence system, while morphological analyses and cytokine quantification were performed using phase-contrast microscopy and multiplex immunoassays. Treatment of macaques with THC protected the astrocytes from virus-induced senescence. Further, THC facilitated a rapid recovery from Tat-induced decline in astrocyte adhesion, suggesting a compensatory effect. THC promoted glial process elongation and morphological complexity, indicative of a shift toward a neuroprotective phenotype. Furthermore, THC significantly reduced inflammatory cytokine secretion, including TNF-α, IL-6, and IL-1β, in an apparently dose-dependent manner. These findings suggest that THC may modulate neuroinflammation in PLWH by promoting astrocytic survival, suppressing inflammatory cytokine secretion, and enhancing neurotrophic signaling. However, prolonged exposure to high-dose THC may negatively impact glial survival. The results underscore the complexity of cannabinoid signaling in the CNS and highlight the potential of cannabinoid-based interventions to mitigate HIV-associated neuroinflammation.

## INTRODUCTION

### HIV-Associated Brain Injury: Persistent Neuroinflammation Despite Viral Suppression

The advent of combination antiretroviral therapy (cART) has greatly improved the life expectancy of people living with HIV (PLWH), yet the prevalence of HIV-Associated Brain Injury (HABI) has remained around 50%. HABI encompasses a spectrum of cognitive impairments resulting from HIV infection and associated neuroinflammation. Unlike prior classifications focused solely on neurocognitive decline, HABI accounts for the broader spectrum of brain injuries influenced by HIV, including neuroinflammatory cascades, glial dysfunction, and vascular contributions to cognitive impairment (Nightingale et al. 2023). Proposed mechanisms of HABI include chronic neuroinflammation driven by HIV viral reservoirs, persistent immune activation despite viral suppression, and endothelial dysfunction contributing to blood-brain barrier (BBB) impairment (Van Zandt and MacLean 2023). Infected microglia and astrocytes serve as key mediators of neuroinflammation, releasing proinflammatory cytokines and extracellular vesicles (EVs) that propagate neural damage (Horn and MacLean 2021). Additionally, the molecular and cellular mechanisms underlying chronic inflammation in glia are not well understood, limiting the development of targeted therapeutics. The persistence of HABI in virally suppressed individuals underscores the challenge of addressing HIV-induced neuroinflammation. This may be indicative of an earlier onset of otherwise normal brain aging processes or highlight a need for tools more specifically designed to assess HIV-induced cognitive changes.

### The Endocannabinoid System in HIV

Cannabis and cannabinoids are the most documented drug of abuse (DoA) in PLWH (Shiau et al. 2017). Both prescribed and self-administered, they are used for alleviation of symptoms associated with disease progression such as neuropathic pain management and appetite stimulation. The primary active components of Cannabis sativa, Δ9-tetrahydracannabinol (Δ9-THC) and cannabidiol (CBD), bind to endogenous cannabinoid receptors (endoCBRs) with varied affinity. The endocannabinoid system is comprised of endoCBRs (CB1 and CB2), their interacting lipids, endogenous cannabinoids, and the metabolic enzymes responsible for their formation and degradation (Piomelli 2003).

EndoCBRs act as presynaptic modulators of inhibitory and excitatory neurotransmitters within the CNS and beyond (Alger 2002; Wilson and Nicoll 2001; Pandey et al. 2009). Found in neurons and endothelial cells, CB1 is one of the most abundant G-protein-coupled receptors (GPCR) expressed in the brain (Howlett and Abood 2017). Activation of CB1Rs in neurons reduces presynaptic GABA release, eliminates GABAergic inhibitory control of postsynaptic neurons, and excites these postsynaptic neurons through dis-inhibition (Alger 2002). Through the binding of macrophage/microglia/astrocyte-expressed CB2, a cascade of GCPR events exerts immunomodulatory effect on target cells by inhibiting cytokine and presynaptic neurotransmitter release that ultimately alters neighboring extracellular environments. EndoCBs act through CB2 to inhibit IL-12 and IL-23 production in microglia and enhance IL-10 production by NF-_K_B suppression, highlighting the importance of both downregulating proinflammatory pathways and stimulating anti-inflammatory cytokine release (Correa et al. 2009; 2010).

### Cannabinoid Modulation of HIV Neuroinflammation

The expression and function of endocannabinoid receptors is altered upon HIV infection even in the absence of exogenous cannabinoid intake (Yadav-Samudrala et al. 2024; Smyth and Collister 2025). In cases of HIV encephalitis, an upregulation of CB1 was observed in neurons and microglia as well as an upregulation of CB2 in microglia, perivascular macrophages, and astrocytes leading to higher cell activation frequency (Cosenza-Nashat et al. 2011). PET imaging in virally suppressed patients demonstrates that the number of activated glia in the CNS of PLWH is inversely correlated with cognitive performance in virally suppressed patients, supporting this proinflammatory hypothesis (Rubin et al. 2018). Putative mechanisms behind this feed-forward loop of inflammation include prolonged cART toxicity, persistent viral reservoirs within the CNS, and damage associated with the BBB (Starr, Jordan-Sciutto, and Mironets 2021).

Just as HIV infection alters the endocannabinoid system, cannabinoids have been shown to alter the dynamics of HIV infection, as cannabis exposure correlates with a lower incidence of HABI (Caitlin Wei-Ming Watson et al. 2020; 2023). Cannabis-using PLWH display equal or lower viral load and lower circulating HIV nucleic acid concentrations than non-users (Thames et al. 2016; Bredt et al. 2002; Chaillon et al. 2020). Cannabis use is also linked to increased CD4+ and CD8+ T-cell counts (Thames et al. 2016; Keen et al. 2019), fewer markers of immune activation (Chaillon et al. 2020; Manuzak et al. 2018), and decreased levels of inflammatory cytokines (C. Wei-Ming Watson et al. 2021; Kumar et al. 2019).

We hypothesized that SIV infection would induce senescence and dysfunction in glial cells, but that these may be inhibited by cannabinoid treatment. Our experiments examined the extent to which treatment of macaques with THC could reverse senescence in astrocytes induced by SIV infection. Further, we sought to determine if THC could reverse this senescence when applied to senescent cultures or reduce the release of pro-inflammatory cytokines. Our novel findings are important for future translational studies in animal models and serve as early insights to the prevention of HABI, and glial contributions to the inflammatory environment within brain in the context of SIV infection managed by long-term ART and how this is affected by cannabinoid administration.

## MATERIALS & METHODS

### Ethical Considerations

All procedures involving rhesus macaques were approved by the Tulane Institutional Animal Care and Use Committee (IACUC) and adhered to the standards set by the Association for Assessment and Accreditation of Laboratory Animal Care (AAALAC) and the Guide for the Care and Use of Laboratory Animals (National Research Council, National Academies Press, Washington, DC). The use of HIV Tat protein was conducted in accordance with NIH guidelines for biosafety and biosecurity.

### Cell Culture

For these studies, cell cultures were obtained from three populations of macaques: SIV-negative, SIV-infected on ART, and SIV-infected treated with THC. Primary mixed glial cultures were derived from the frontal lobes of rhesus macaques (*Macaca mulatta*) following necropsy. Using previously established protocols (Guillemin et al. 1997; Andrew G. MacLean et al. 2002; A. G. MacLean et al. 2004; Andrew G. MacLean et al. 2004; Renner et al. 2013), meninges were carefully removed, and cortical tissue was finely diced with sterile scalpels. The resulting tissue fragments were enzymatically digested with 0.25% trypsin (Invitrogen, Carlsbad, CA) and DNAse (4 U/ml, Worthington, Lakewood, NJ) at 37°C for 60 minutes, followed by trituration and filtration through 110 μm pore filters (Sigma). The cell-rich slurry was subjected to three rounds of centrifugation at 1,000 rpm, washed, and resuspended in M199 medium supplemented with 0.7mM sodium bicarbonate and 5% fetal bovine serum. Cells were initially cultured in T-25 flasks, with media replenished at 24 hours.

### Growth and Morphology Analysis

For clarity, the methods for these studies are divided into three sections:

1: Cultures from THC treated, SIV-infected animals (animals received THC),
2: Cultures from SIV-infected animals (cell cultures received THC), and
3: Cultures from naïve animals.

1: Cultures from THC treated, SIV-infected animals (animals received THC): Primary glial cultures from SIV-infected animals that were administered THC for at least 90 days were utilized to assess differences in cell growth and glial morphology. After 24 hours in culture, the cells were gently washed and fresh media added. Media was then replaced twice weekly. Phase-contrast images were taken weekly using the 10x objective of a Zeiss Axiovert 100 microscope from 10 non-overlapping fields for each flask.
2: Cultures from SIV-infected animals (cell cultures received THC): Primary glial cultures from SIV-infected animals on a stable regimen of ART were utilized to assess differences in cell growth and glial morphology with *in vitro* exposure to THC. Cultures were washed to remove debris at 24 hours, and then beginning at 48 hours media was collected and replaced with fresh media or media containing 5 µM, or 10 µM THC that was coded to maintain blinding for image collection and analysis. Media collections and replacements were then performed every 48 hours to ensure continued exposure to THC. Phase-contrast images were taken using the 10x objective of a Zeiss Axiovert 100 microscope from 10 non-overlapping fields for each flask prior to application of THC, at 24hrs post first application, then every 72 hrs. Cell counts and branching complexity were quantified using FIJI ImageJ software.
3: Cultures from naïve animals. After 24 hours in culture, the cells were gently washed and fresh media added. Media was then replaced twice weekly until approximately 80% confluent. Cells were subcultured into T-75 flasks or seeded onto 16-well xCELLigence E-Plates (Agilent) for real-time analysis of adhesion and proliferation. Cells were allowed to adhere for at least 24 hours before experimental treatments were introduced, ensuring stable attachments and baseline impedance measurements. Morphological assessments were used to determine whether cannabinoids promoted astrocytic stellation or counteracted Tat-induced cellular activation.

### xCELLigence Assay: real time measures of cell adhesion, proliferation, and morphology change

Cell adhesion and proliferation was measured and plotted automatically using an xCELLigence RTCA analyzer as previously described (Renner et al. 2013; Andrew G. MacLean et al. 2014). Each E-Plate was equilibrated by adding 50uL of complete media to each well and incubating for 15 minutes at 37°C in 5% [CO_2_] before background impedance was established. 20,000 cells (from SIV-negative animals) suspended in 100uL of complete media were applied to each well with measurements taken at 5-minute intervals. Experimental conditions were performed in quadruplicate as recommended in the technical manual of the RTCA instrument. 24 hours after plating, wells were treated with control media or HIV Tat (NIH HIV Reagent Program, Division of AIDS, NIAID, NIH). Following continued Tat exposure, cells were treated with 5uM THC (The National Institute on Drug Abuse (NIDA) Drug Supply Program (DSP)). Traces were plotted and analyzed using the installed software.

### Cytokine Measurements

Media was collected from flasks at each media change, frozen at –80°C and analyzed for the presence of senescence-associated secretory phenotype (SASP) markers using an eleven-plex immunoassay designed for non-human primates (Assay Genie, Ireland (Csiszar et al. 2012)). Cytokines were measured on an LSRFortessa (BD Biosciences) and raw FCS files analyzed by Assay Genie.

To determine whether THC treatment influenced neurotrophic signaling, fibroblast growth factor basic (FGF-B) levels were measured in culture media using a single-plex immunoassay. FGF-B is a crucial mediator of synaptic plasticity and neuroprotection, and its upregulation would suggest a potential mechanism by which THC exerts its neuroprotective effects.

### Statistical Analysis

Data were analyzed using GraphPad Prism software. Comparisons between experimental conditions were conducted using one-way ANOVA followed by Tukey’s post-hoc test for multiple comparisons. Results were expressed as mean ± standard error of the mean (SEM), and statistical significance was defined as *p* < 0.05.

## RESULTS

### THC Prevents Senescence in Glial Cultures from SIV-Infected Macaques

In cultures derived from research naïve animals, small cell colonies emerged between days 7– 10, with cells adopting a spindle-like or polygonal astrocytic morphology (Figure 1). By day 14, these control cultures formed multiple areas with continuous monolayers characteristic of mature astrocyte populations, with stratification and cell stacking over time.

**Figure 1.**
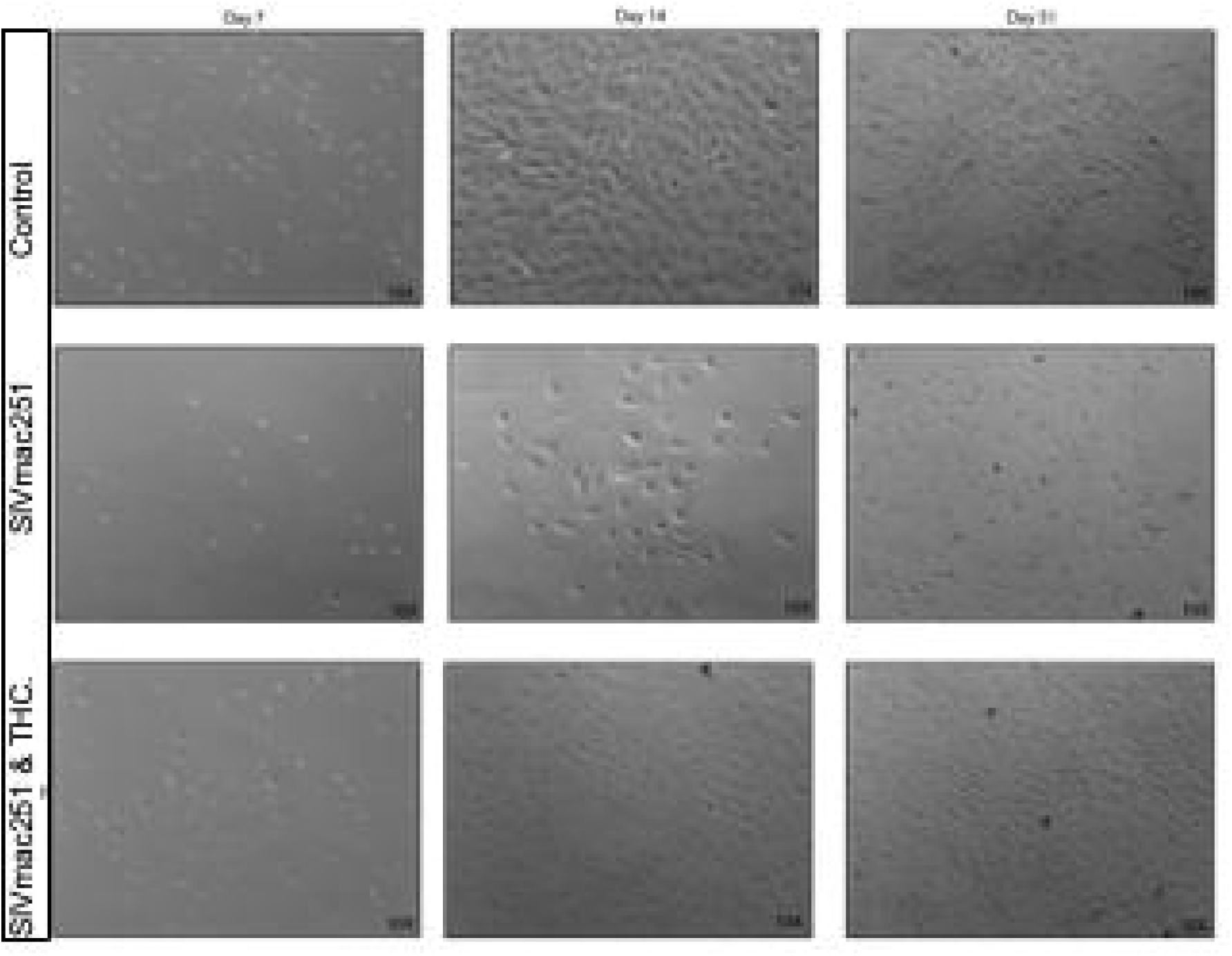
THC Prevents Senescence in Glial Cultures from SIV-Infected Macaques. Representative phase-contrast microscopy images of primary mixed glial cultures derived from naïve rhesus macaques compared to SIV-infected, and SIV-infected treated with THC. Control cultures from naïve young to middle-aged macaques typically form small colonies between days 7–10. By day 14, cells within these colonies reach confluence, forming a uniform monolayer that continues to proliferate and stratify under healthy conditions. Glial cultures from SIV-infected animals displayed reduced cell adhesion at early time points, with fewer colonies forming by day 7. By day 14, these cultures demonstrated lower cell density and increased morphological heterogeneity, with astrocytes displaying a more reactive, hypertrophic phenotype. In contrast, cultures from SIV-infected animals that had been treated with THC while alive grew, albeit at a slightly lower rate compared with control cultures.

In contrast, cultures from SIV-infected animals exhibited a near complete lack of proliferation over the first 14 days and fail to form colonies or “islands” of cells that are apparently necessary for successful growth (unpublished observation). By day 7 (not shown), fewer cells remained adhered to the culture flasks, and colony formation was notably diminished. By day 14, these cultures displayed lower overall cell density, with hypertrophic astrocytes exhibiting enlarged somas and retracted processes, indicative of a reactive phenotype. By day 51, glia from SIV-infected animals exhibited persistent morphological abnormalities, including cellular debris, disrupted monolayer formation, and increased heterogeneity in morphology, suggesting a shift toward a neurotoxic, pro-inflammatory state.

Cultures derived from SIV-infected, THC-treated animals, by contrast, were able to adhere, form colonies, and proliferate, although at a rate slower than those from research naïve animals. It was noted that there may have been less stratification, or piling up, of the cultures as cells approached confluence.

With this context established, we next investigated the extent to which HIV Tat protein in isolation may be responsible for changes in astrocyte adhesion, morphology, and cytokine release and if these can be reversed or inhibited with cannabinoid treatment.

### Tat Disrupts Astrocyte Adhesion and Morphology

To assess the effects of Tat on astrocytes, we measured adhesion of mixed glial cultures in real time over a 40-hour period following exposure to Tat (2μM, 3μM, or 5μM). All data are baselined to vehicle-containing media, with impedance graphed as cell index (CI) using proprietary software from Agilent. The 3μM Tat group demonstrated a downward trend, though not significant, while the low tat (2μM) group remained close to baseline, indicating a minimal impairment. However, 5μM Tat induced a rapid decrease in CI, which remained low for 48 hours (Fig 2A), indicating a marked reduction in cell adhesion. Replacement of the Tat-containing media with control media allowed a recovery of the CI (not shown), indicating minimal loss in viability.

**Figure 2.**
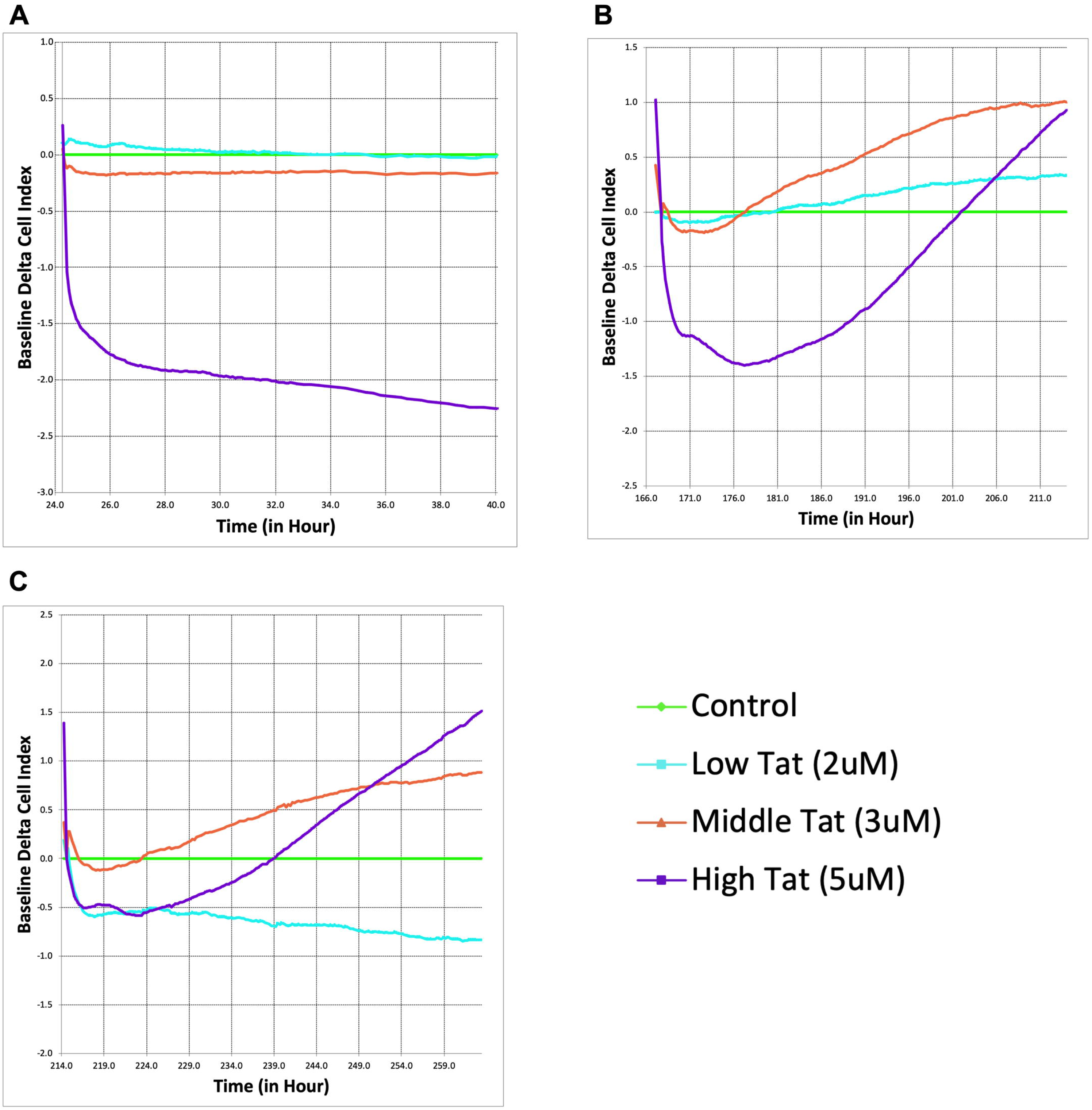
THC Counteracts Tat-Induced Astrocyte Dysfunction and Enhances Cellular Recovery. Real-time xCELLigence assay measuring the baseline delta cell index of astrocytes exposed to HIV Tat (2μM, 3μM, and 5μM). Higher Tat concentrations resulted in a significant, dose-dependent decline in astrocyte adhesion and viability over time, as indicated by a decrease in cell index (A). Following THC administration (5μM), cell index recovery was observed, particularly in the higher Tat groups (B). Indeed, the cell indices rebounded beyond the glial cultures that did not receive Tat, indicating a pronounced compensatory response. A second THC administration further enhanced astrocyte recovery, with the higher Tat groups displaying a sustained increase in cell index. Curiously, the cells treated with the lowest Tat concentration (2µM) had a sustained decrease in the cell index, even in the presence of THC.

### THC Counteracts Tat-Induced Astrocyte Dysfunction and Enhances Cellular Recovery

To determine whether THC counteracts Tat-induced astrocyte dysfunction, cells were administered a single 5μM dose of THC following five days of Tat exposure (Figure 2B). Surprisingly, rather than stabilizing at control levels, the 5μM Tat group exhibited a dramatic increase in CI, far exceeding the control baseline. This suggests that THC induces an exaggerated compensatory response in stressed astrocytes, potentially through cytoskeletal remodeling, increased adhesion, and / or enhanced metabolic activity. Meanwhile, the 3µM Tat group displayed a more moderate recovery, surpassing the control baseline, whereas cells treated with 2µM Tat group showed only a very slight deviation from control cells.

Following a second THC administration (Figure 2C), the CI decreased to a lesser degree and responded more rapidly than previously. The 3μM Tat treatment demonstrated an almost identical response to the first THC treatment, while the low Tat group remained relatively unchanged, or even decreased. These suggest that the impact of THC is most pronounced in astrocytes subjected to more severe Tat-induced stress.

### Exogenous THC Fails to Reverse Glial Senescence in Cultures from SIV-infected Macaques

Given the dual role of THC in both promoting trophic support and inducing cellular stress, we examined its impact on glial survival over time. Primary mixed glia from SIV-infected macaques were cultured in control medium, or with either 5μM or 10μM THC, and viability assessed by counting the number of cells (Figure 3). Contrary to previous reports suggesting trophic effects of cannabinoids, our data reveal a progressive decline in glial cell viability with a pronounced reduction in cell count when treated with 10µM THC, and essentially no deleterious effect at 5µM.

**Figure 3.**
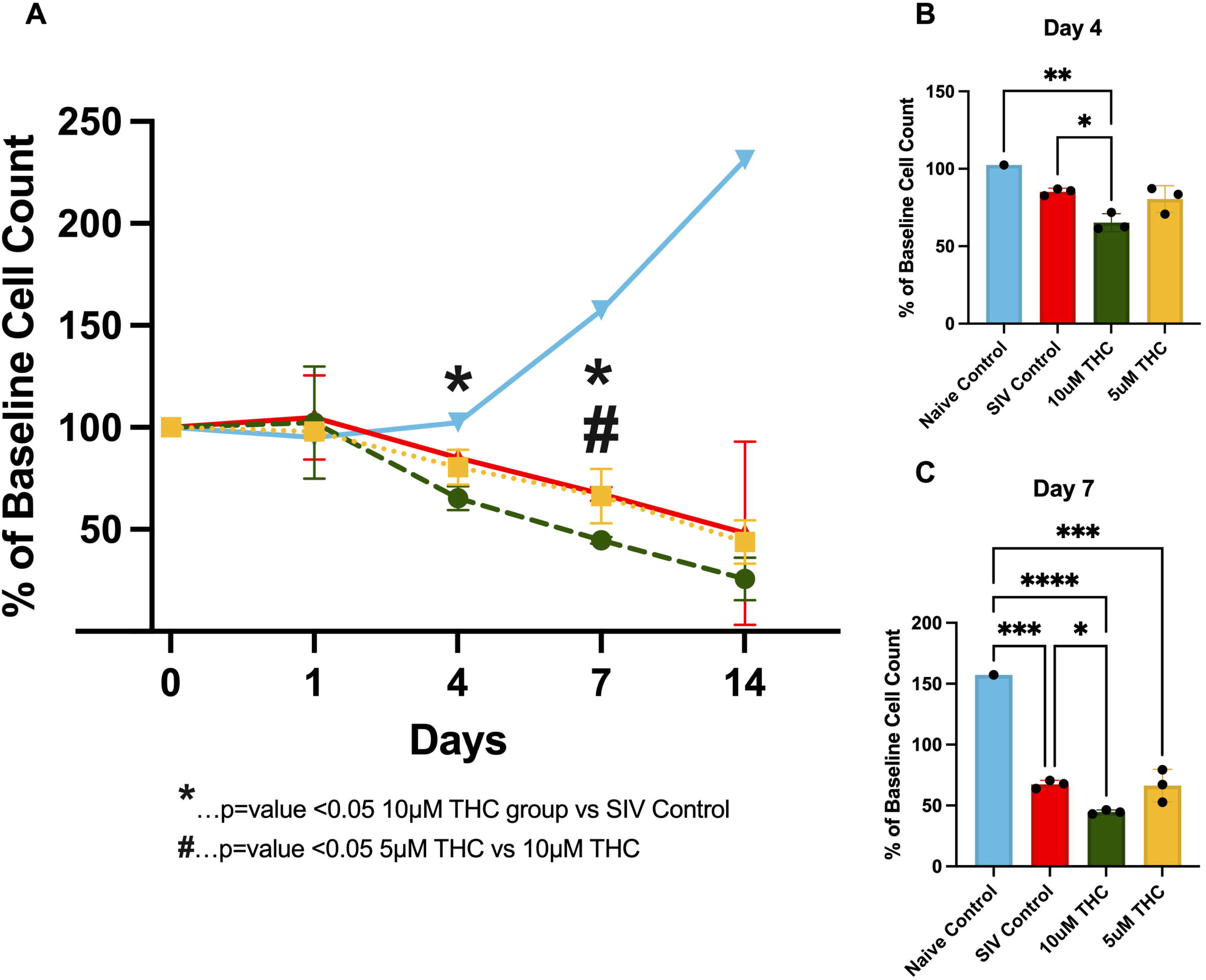
Exogenous THC Fails to Reverse Glial Senescence in Cultures from SIV-infected Macaques. Quantification of astrocyte viability over initial 14 days in cultures from animals infected with SIV (A). THC exposure resulted in a progressive decline in astrocyte survival, particularly at the 10μM concentration. By Day 4 in culture, a significant reduction in cell count was observed in the 10µM group compared to control (p < 0.05). By Day 7, 10μM THC resulted in a significantly greater decline in astrocyte viability relative to 5μM THC and control (p < 0.05), suggesting a dose-dependent cytotoxic effect. Data are expressed as % baseline cell count ± SEM; statistical significance determined by one-way ANOVA followed by Tukey’s post hoc test.

By Day 4 (Figure 3B), 10µM THC-treated groups showed a significant reduction in cell count compared to the control. This decline became more pronounced by Day 7 (Figure 3C), where 10µM THC-treated cultures were significantly lower than the 5μM and control groups. These findings suggest that while cannabinoids modulate glial function, chronic exposure to higher THC concentrations may exert detrimental effects on glial viability, highlighting the importance of dose considerations when evaluating THC’s therapeutic potential in neuroinflammatory conditions such as HIV-associated neuroinflammation (Cosenza-Nashat et al. 2011).

### THC Enhances Glial Process Formation and Morphological Complexity

To determine if this is truly fewer cells, or fewer cells relative to the untreated control, we analyzed the development of processes on the glia when treated with THC starting immediately after plating the cells (Figure 4). A clear dose-dependent enhancement in glial process formation was observed, with the 10 μM THC-treated cells exhibiting greater process length and branching complexity than the 5 μM-treated and control groups.

**Figure 4.**
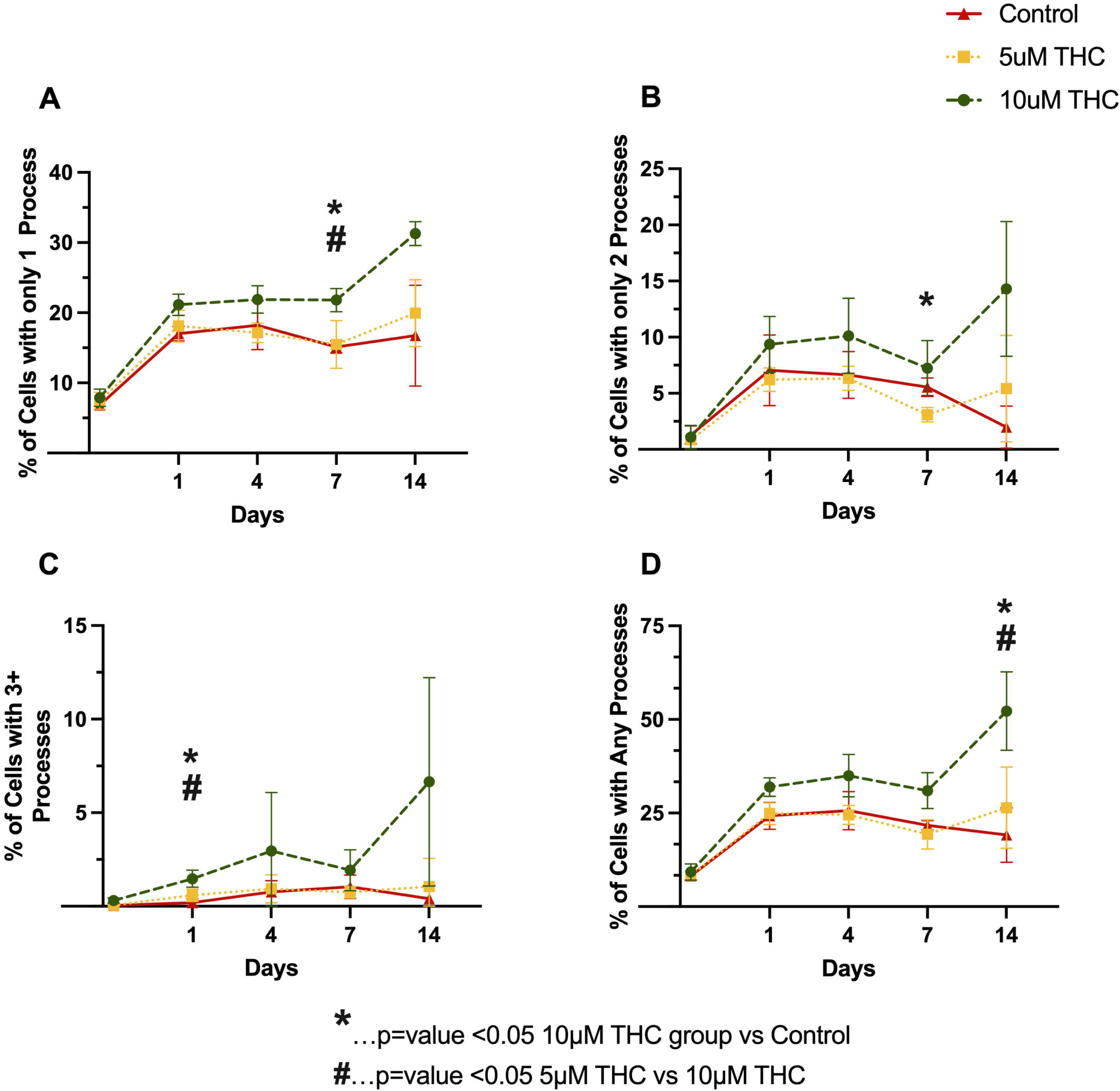
THC Enhances Glial Process Formation and Morphological Complexity. Quantification of glial morphology following THC exposure, assessing the percentage of cells with only one process (A), only two processes (B), three or more processes (C), and any identifiable processes (D) over the initial 14-days in culture. THC-treated astrocytes exhibited increased process formation in a dose-dependent manner, with 10 μM THC-treated cells displaying greater process elongation and branching complexity relative to controls. By Day 7 and Day 14, 10 μM THC significantly increased the percentage of cells with elongated processes compared to 5 μM THC and control (p < 0.05). Data are presented as mean ± SEM; statistical significance determined using one-way ANOVA.

### THC Suppresses Senescence-Associated Secretory Phenotype in Mixed Glia Cultures

To determine if the glial polarization and adhesion altered secretion of pro-inflammatory and pro-senescence cytokines, we employed an 11-plex immunoassay of glial cultures derived from SIV-infected animals. Levels of IL-6 and MCP-1 were significantly reduced by treatment with either 5µM or 10µM THC by ten days in culture (Figure 5), whereas, 10µM THC-treated cultures had significantly lower levels of IL-1B, INF-γ, and IL-8. The other cytokines measured showed no significant alteration. The suppression of senescence-associated proinflammatory cytokine secretion is particularly relevant to HIV-associated neurocognitive disorders, as chronic neuroinflammation contributes to neuronal damage, synaptic dysfunction, and cognitive decline. These data reinforce previous findings that CB2 receptor activation dampens neuroinflammatory pathways, thereby alleviating immune activation and excitotoxic stress within the CNS.

**Figure 5.**
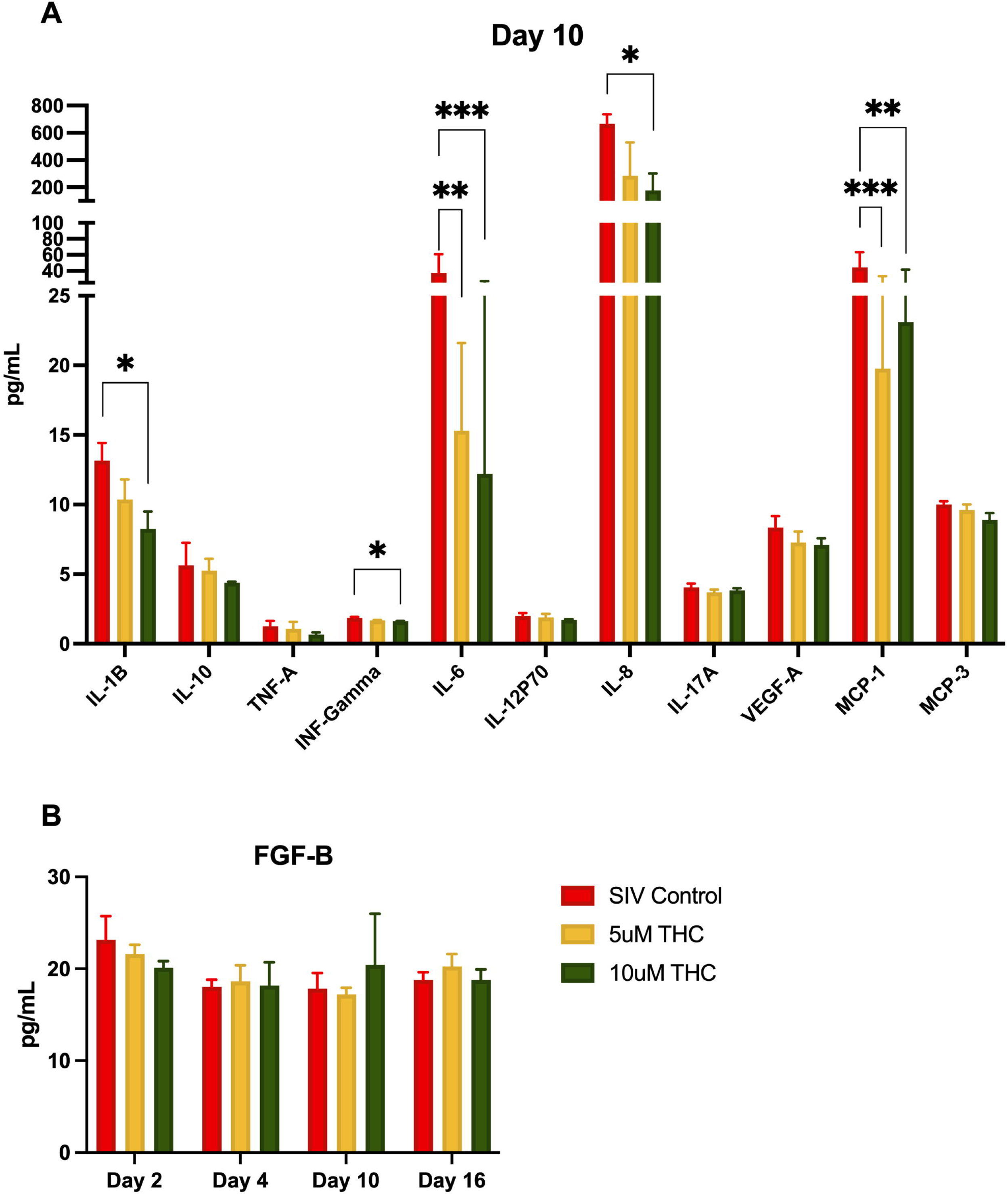
THC Suppresses Senescence-Associated Secretory Phenotype in Mixed Glia Cultures. Treatment of primary glial cultures from SIV-infected macaques with THC altered secretion of multiple cytokines (A), with IL-1Β, INF-γ, and IL-8 significantly reduced by 10µM THC (*p < 0.05, **p < 0.01, ***p < 0.001). Both 5µM and 10µM THC reduced secretion of IL-6 and MCP-1. While other cytokines had reduced expression (including TNF-a, this did not reach the significance threshold). Fibroblast growth factor-basic (FGF-B) levels were quantified over time, revealing a slight, though not significant, increase in FGF-B secretion in 10µM THC-treated astrocytes (B). Data are presented as mean ± SEM; statistical significance determined using one-way ANOVA followed by Tukey’s post hoc test.

FGF-B plays a crucial role in neuroprotection, synaptic plasticity, and glial homeostasis, and its significant upregulation in the serum of HIV-infected individuals reinforces THC’s potential anti-inflammatory effects (Ray et al. 1999; Ascherl et al. 2001). Therefore, we measured release of FGF-B in the same culture supernatant as above. We were somewhat surprised that there was no significant change in the level of FGF-B with either THC treatment (B), although there was a minor increase at ten days of treatment with 10µM THC.

## DISCUSSION

In this study, we provide new insights into how SIV infection and HIV Tat protein affect glial function, morphology, and survival, with a particular focus on the potential modulatory role of cannabinoids in HIV-associated neuroinflammation. By integrating real-time impedance assessments, morphological analysis, cytokine profiling, and neurotrophic factor measurements, we demonstrate both the detrimental effects of viral neurotoxins and the complex role of THC in regulating glial responses.

Before investigating the effects of Tat and THC, we first examined how SIV infection itself alters glial proliferation and morphology (Figure 1). In agreement with previous *in vivo* studies demonstrating glial dysfunction and neuroinflammation in HIV/SIV infection, SIV+ glial cultures exhibited reduced adhesion, delayed colony formation, and disrupted monolayer development. By later time points, SIV+ cultures showed persistent abnormalities, including hypertrophic morphologies, with enlarged somas and retracted processes, indicating a shift toward a reactive phenotype that may exacerbate neuroinflammatory signaling. Notably, astrocytes derived from SIV+ animals that had received THC, had morphologic phenotypes more similar to those grown from naïve animals, further supporting the hypothesis that SIV infection primes glial populations for dysfunction, but that these may be inhibited by cannabinoid treatment.

Given this pre-existing glial dysfunction in SIV+ cultures, we next investigated whether HIV Tat exposure alone can induce these impairments and whether THC has a compensatory or neuroprotective effect. Tat exposure induced a dose-dependent decline in glial cell index, demonstrating its well-documented inflammatory / negative effects on neural cells (Cosenza-Nashat et al. 2011; Rubin et al. 2018). We were surprised to find that treatment with 5μM THC not only reversed the Tat-induced decline in cell index but, in two of the three Tat exposure groups, led to cell index values exceeding those of untreated controls (Figure 2). These suggest that Tat and THC may interact in a concentration-dependent manner, with higher Tat exposure shifting glial cells into a state more receptive to cannabinoid-mediated neuroprotection. The ability of THC to not only prevent further decline but enhance cell index in the presence of high Tat concentrations may indicate a potential compensatory or trophic effect of cannabinoid signaling in the context of Tat-induced neuroinflammation. Interestingly, repeated THC administration produced a sustained recovery in cell index, particularly in the high Tat group, raising the possibility of a compensatory response in surviving astrocytes. While Tat is traditionally regarded as neurotoxic, these findings suggest that Tat-induced glial activation may create a therapeutic window—in which cannabinoid receptor signaling could exert a compensatory effect. This was unexpected, especially considering that in glia derived from SIV-infected animals, THC not only failed to restore baseline cell counts but worsened the decline (Figure 3).

The decline in cell numbers (Figure 3) highlights the need for further investigation into whether the increased cell index reflects improved astrocyte function or a maladaptive response, such as excessive glial activation or metabolic stress. Understanding how Tat primes astrocytes for a heightened response to THC could provide valuable insight into the dynamics of neuroimmune signaling in PLWH. Future studies will need to investigate whether the increased cell index in the high Tat group reflects improved astrocyte function or represents a pathological overcompensation, such as excessive glial activation or metabolic dysregulation (League et al. 2024; Ton and Xiong 2013; Borgmann and Ghorpade 2015; Fan and He 2016; Hermes et al. 2021; Cotto, Natarajanseenivasan, and Langford 2019).

Further supporting THC’s role in neuroprotection or possibly a reparative astrocytic phenotype, THC modulated glial morphology, promoting stellation and increased process complexity in primary mixed glial cultures (Figure 4). Given that astrocyte morphology is closely linked to functional outcomes in neuroinflammatory conditions, these findings suggest a shift toward a neuroprotective phenotype (Renner et al. 2012, Lee et al. 2014). The changes in astrocyte morphology observed in this study are consistent with cannabinoid receptor activation attenuating glial reactivity and promoting cellular adaptations that support neuronal survival. Similar effects have been observed in models of neurodegenerative disease, where cannabinoid treatment modulates glial activation states and reduces inflammatory signaling (Henriquez et al. 2020).

CB2 receptor activation enhances anti-inflammatory cytokine production, including IL-10, while suppressing proinflammatory mediators such as IL-12 and IL-23 (Correa et al. 2010). Our study adds to this by demonstrating that THC significantly reduced the secretion of senescence-associated proinflammatory cytokines, including IFN-γ, IL-6, and IL-1β, reinforcing its potential to counteract chronic neuroinflammation (Csiszar et al. 2012). Cannabis use in HIV-positive individuals is associated with lower levels of CNS inflammation and a reduced risk of neurocognitive impairment (Caitlin Wei-Ming Watson et al. 2020; 2023). Taken together, these findings suggest that THC fosters an anti-inflammatory, neuroprotective environment by modulating glial morphology, cytokine secretion, and trophic factor production, even though it may not enhance overall glial cell survival. This aligns with prior studies indicating that CB1 and CB2 receptor activation can suppress neuroinflammatory pathways and influence glial function (Correa et al. 2009; Manuzak et al. 2018). Given that cannabinoids regulate immune activation and synaptic function, the results presented here suggest that THC exerts neuroprotective and anti-inflammatory effects to counteract key mechanisms underlying HIV pathogenesis through modulation of glial adhesion, proliferation, and cytokine secretion.

Given the persistent burden of HABI despite effective antiretroviral therapy, novel strategies targeting neuroinflammation and synaptic dysfunction are urgently needed. These current findings provide compelling evidence for cannabinoid-based therapeutics as a strategy to mitigate neuroinflammation and synaptic dysfunction in PLWH. However, they also underscore the complexity of cannabinoid signaling in the CNS, emphasizing the need for precise modulation of the endocannabinoid system to achieve optimal neuroprotective outcomes. THC’s ability to suppress inflammatory cytokines, promote glial homeostasis, and enhance neurotrophic signaling suggests that cannabinoid-based interventions may offer a viable approach to slowing HAND progression, though the impact on overall glial viability remains a critical consideration.

An important consideration is the differential effects of cannabinoid compounds. While this study focuses on Δ9-THC, cannabidiol (CBD) has also been shown to exert anti-inflammatory and neuroprotective effects without the psychoactive properties of THC (DeMarino et al. 2022; Kaddour et al. 2022). Comparative analyses of THC and CBD in modulating cognitive function, neuroinflammation, and neurodegeneration in SIV-infected macaques may provide valuable insights into the development of targeted cannabinoid therapies. Further research using *in vivo* models and clinical studies is needed to assess the long-term efficacy and safety of THC in preserving neurocognitive function in people living with HIV.

This study provides compelling evidence that THC exerts neuroprotective and anti-inflammatory effects in HIV-associated brain injury (League et al. 2024) and supports the growing interest in cannabinoid-based therapeutics for neuroinflammatory disorders (Jana et al. 2024; Möller et al. 2025; Cárdenas-Rodríguez et al. 2024; Alraddadi et al. 2025). Understanding the molecular pathways underlying this response will be critical for developing targeted cannabinoid-based interventions for HIV-associated neuroinflammation. Future research will focus on elucidating the precise molecular mechanisms underlying THC’s effects on glial function and assessing the long-term efficacy and safety of cannabinoids in PLWH.

## ACKNOWLEDGEMENTS

Funding Statement: NHP studies in the MacLean Lab are supported by NIH funds: R21-MH113517, R01-HL152804, R01-NS104016, R21-MH125716, U42OD024282, U42OD010568, and the TNPRC base grant P51-OD11104. Flow Cytometry data acquisition and analysis was supported by NIH S10 OD026800. Tulane University also supports this research through the Tulane Brain Institute and the Tulane Neuroscience Program.

Antiretroviral drugs were generously provided by Gilead Sciences, Inc. (TDF and FTC) and ViiV Healthcare (DTG).

Simian Immunodeficiency Virus (SIV)mac239 Tat Region, ARP-12765, contributed by DAIDS/NIAID, was obtained through the NIH HIV Reagent Program, Division of AIDS, NIAID, NIH.

5mg/mL Δ9-Tetrahydrocannabinol (THC) in ethyl alcohol (DEA code 7370) was provided the National Institute on Drug Abuse (NIDA) Drug Supply Program (DSP).

Fresh brain tissue from several NIH-funded studies were made available at necropsy. We are indebted to the several collaborators over the years, including Drs Ronald Veazey, Nicholas Maness, Mahesh Mohan, and Andrew Lackner. We would like to thank the pathologists and staff of the Anatomic Pathology Core RRID: SCR_024606, Clinical Pathology Core RRID: SCR_024609, Confocal Microscopy and Molecular Pathology Core RRID: SCR_024613, Virus Characterization, Isolation, Production and Sequencing Core RRID:SCR_024679, and Flow Cytometry Core RRID: SCR_024611, SCR_008167. This work could not happen without the dedicated staff and veterinarians of the Division of Veterinary Medicine.

## FIGURE LEGENDS

**Supplemental Figure 1.** Representative images of SIV+ mixed glial cultures treated with THC. Phase-contrast microscopy images of primary mixed glial cultures derived from SIV+ rhesus macaques following administration of 5μM or 10μM THC, compared to control conditions, over a 14-day period. Images were captured every 48 hours to assess changes in cell number, density, morphology, and process formation over time. While the higher concentration of THC led to a progressive reduction in overall cell density (as observed in Figure 3), surviving glial cells exhibited enhanced stellation and process elongation: a possible shift toward a neuroprotective phenotype. This suggests that THC promotes cytoskeletal remodeling and astrocyte activation, potentially facilitating enhanced synaptic support and immune modulation despite an overall reduction in cell viability. Notably, in the 10μM THC condition, glial cells demonstrated the most pronounced process elongation and branching complexity, possibly compensating for the decline in total cell numbers. These findings highlight the dual role of THC in modulating glial survival and function, where higher THC concentrations may induce cellular stress leading to cell loss, yet simultaneously promote a more complex and supportive astrocytic network.

## Notes

### Competing Interest Statement

The authors have declared no competing interest.

