## Supplementary figures and images for "THC Reverses SIV-Induced Senescence in Astrocytes: Possible Compensatory Mechanism Against HIV Associated Brain Injury?"

### Supplemental Figure 1

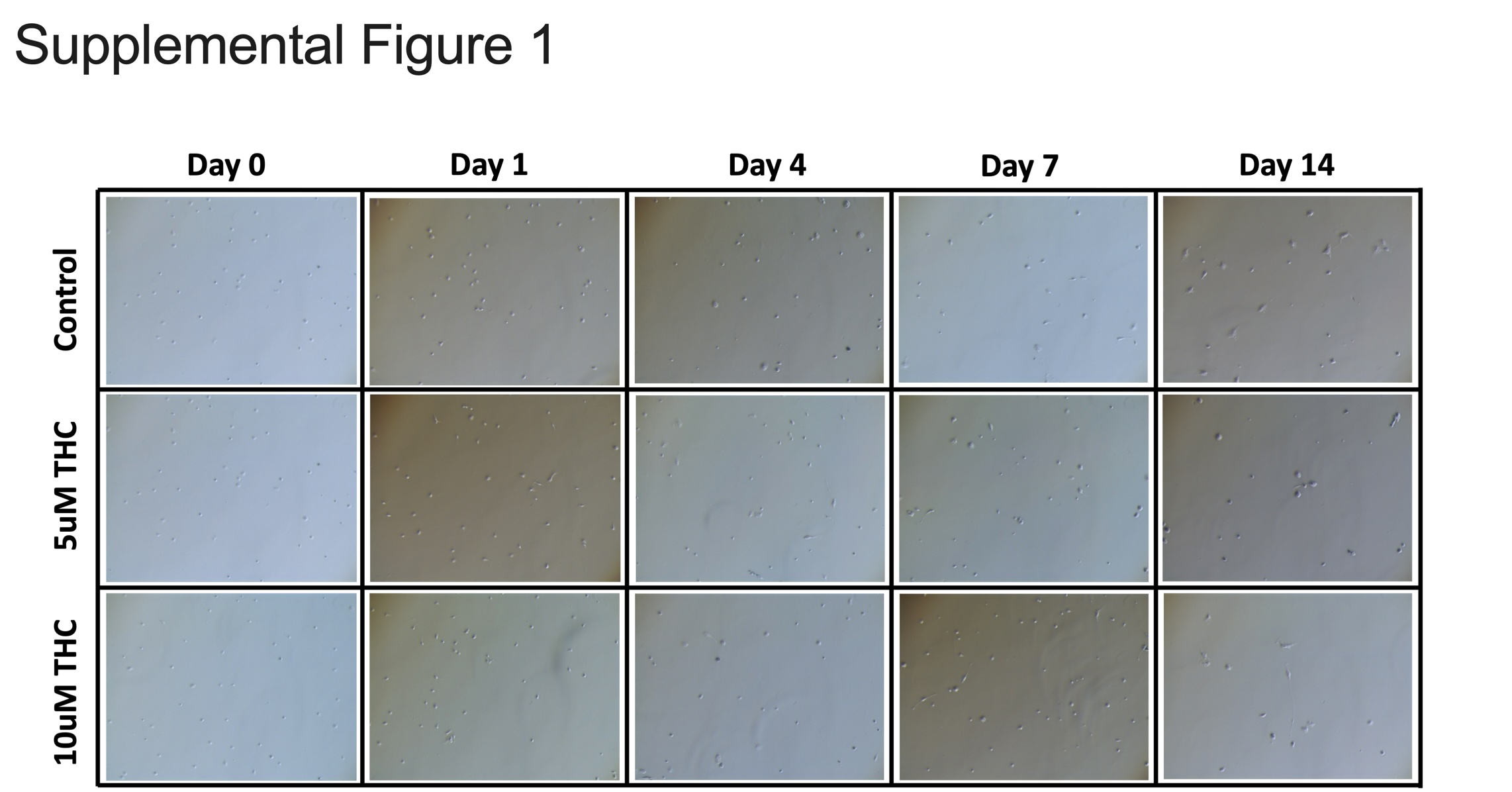
